# Head and Neck Cancer-derived small extracellular vesicles sensitize TRPV1+ neurons to mediate cancer pain

**DOI:** 10.1101/2022.09.06.506411

**Authors:** Kufreobong E. Inyang, Christine M. Evans, Matthew Heussner, Margaret Petroff, Mark Reimers, Paola D. Vermeer, Nathan Tykocki, Joseph K. Folger, Geoffroy Laumet

## Abstract

Severe pain is often experienced by patients with head and neck cancer and is associated with a poor prognosis. Despite its frequency and severity, current treatments fail to adequately control cancer-associated pain, because of our lack of mechanistic understanding. Cancer-derived small extracellular vesicles (Cancer-sEVs) are well-positioned to function as mediators of communication between cancer cells and neurons. Inhibition of Cancer-sEV release attenuated pain in tumor-bearing mice. Injection of purified Cancer-sEVs is sufficient to induce pain hypersensitivity in naïve mice. Cancer-sEVs triggered calcium influx in nociceptors and inhibition or ablation of nociceptors protect against cancer pain. Interrogation of published sequencing data of human sensory neurons exposed to human Cancer-sEVs suggested a stimulation of protein translation in neurons. Induction of translation by Cancer-sEVs was validated in our mouse model and its inhibition alleviated cancer pain in mice. These findings define a role of Cancer-sEVs in cancer pain and identify several druggable targets.

**Graphical abstract:** 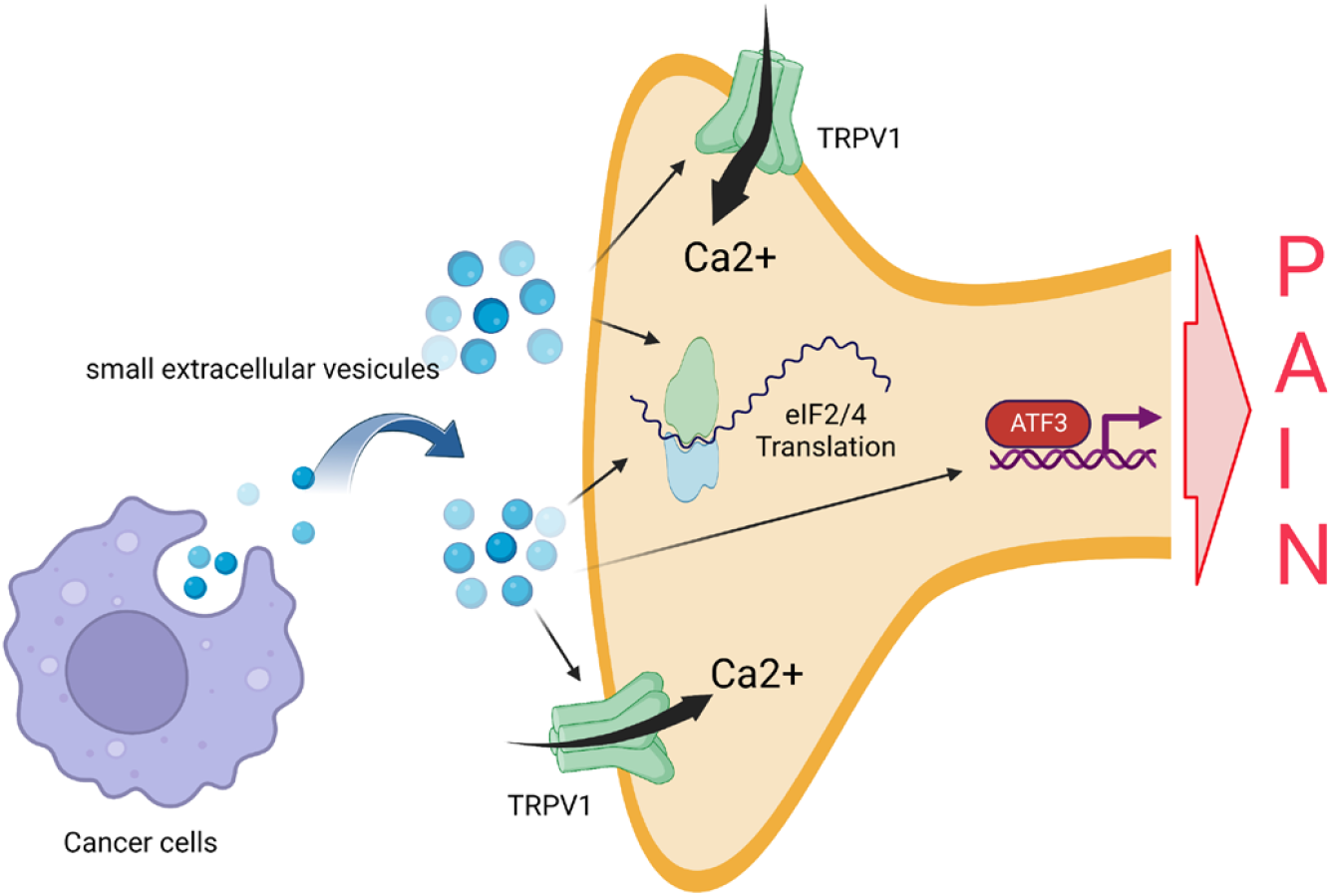

## Introduction

Head and neck squamous cell carcinoma (HNSCC) is the 9^th^ most prevalent form of cancer in the United States (Jemal et al. 2009). Nearly all HNSCC patients experience pain from either the treatment or the cancer itself with approximately 57% reporting tumor-related pain at the time of presentation (Macfarlane et al. 2012, Murphy et al. 2019). In fact, pain is often the main symptom motivating HNSCC patients to seek medical intervention (Marshall et al. 1997, Sato et al. 2011). Patients with HNSCC pain report spontaneous and evoked pain hypersensitivity (Connelly et al. 2004, Salvo et al. 2020). Cancer pain is associated with poor survival outcomes (Herndon et al. 1999, van den Beuken-van Everdingen et al. 2007, Montazeri 2009, Hedberg et al. 2019). Despite the high prevalence and severity of HNSCC-associated pain, current treatments rarely provide adequate pain relief (Connelly et al. 2004, Chen et al. 2010, Lin et al. 2011) because the underlying mechanisms promoting cancer pain are not fully defined. A better understanding of the mechanisms underlying HNSCC pain would facilitate the development of novel non-addictive analgesics and positively impact the quality of life as well as the survival of cancer patients.

While perineural invasion plays a critical role in HNSCC-associated pain (Salvo et al. 2020, Cata et al. 2022), the earliest pain which is experienced before a cancer diagnosis, likely results from secreted factors. HNSCC tumor cells produce and secrete small extracellular vesicles (sEVs; size 30-150 nm) (Whiteside 2017, Wolf-Dennen et al. 2020) which play a role in intercellular communication under normal and pathological settings (Milane et al. 2015, Arenaccio et al. 2017). Once released, sEVs act on local or distant target cells and modulate signaling pathways (Colombo et al. 2014, Ge et al. 2020). sEVs contribute to various processes in cancer progression such as tumor metastasis and immune modulation (Zhang et al. 2015, Zomer et al. 2015, Becker et al. 2016, Kalluri 2016). In addition, HNSCC-derived sEVs induce neurite outgrowth in cultured sensory neurons from humans and mice (Madeo et al. 2018, Amit et al. 2020) and promote tumor innervation by Transient Receptor Potential V1 expressing (TRPV1+) neurons in preclinical models (Madeo et al. 2018). These data indicate that sensory neurons can uptake HNSCC-derived sEVs and change their physiology in response. Although the role of HNSCC-derived sEVs in cancer cell-neuron communication is established, the contribution of sEVs to cancer pain remains unknown.

To model HNSCC in C57Bl/6 wildtype (WT) mice, we injected a previously characterized murine model of human papillomavirus-induced (HPV+) HNSCC called mEERL cells. These cells were derived from oropharyngeal cells isolated from C57Bl/6 mice which were modified to stably express oncogenes. As such, when injected into immunocompetent mice, tumors grow with characteristics that are faithful to the human disease (Williams et al. 2009, Madeo et al. 2018). We previously showed that implantation of mEERL cells in WT mice induces spontaneous and evoked pain (Heussner et al. 2021). Interestingly, in this model there is no neuroinflammation in the spinal cord and blocking IL-1 signaling does not alleviate cancer pain. Similar to human HNSCC, mEERL tumors are innervated by TRPV1+ neurons (Madeo et al. 2018). TRPV1+ neurons are critical for thermal sensitivity and nociception (Caterina et al. 1997). TRPV1 expression is also upregulated in neurons of mouse models of cancer (Asai et al. 2005, Beaudry et al. 2011, Lucido et al. 2019). Therefore, TRPV1+ neurons are an attractive target for treating pain in cancer conditions like HNSCC.

In this study, we explore the contributions of HNSCC-derived sEVs to pain and their communication to TRPV1+ neurons.

## Results

### Blocking cancer-derived sEVs attenuates pain

Given that sensory neurons can take up and respond to HNSCC-derived sEVs (Lopez-Verrilli et al. 2013, Madeo et al. 2018, Amit et al. 2020), we assessed the effects of blocking cancer-derived sEVs on cancer pain. Implantation of mEERL cells into WT mice induced evoked and spontaneous pain (Heussner et al. 2021). Administration of the sEV release inhibitor GW4869 (1.25 mg/kg) attenuates pain hypersensitivity and facial grimacing in tumor-bearing mice (**Figure 1A-C**). Blockage of sEV release is confirmed by the reduction of circulating sEVs (**Figure 1D**). Additionally, implantation of mEERL Rab27a-/+ and Rab27b-/- cells, cancer cells that release a limited amount of sEVs significantly delayed the development of pain hypersensitivity and spontaneous pain (**Figure 1E-G**) compared to parental mEERL cells.

**Figure 1.**
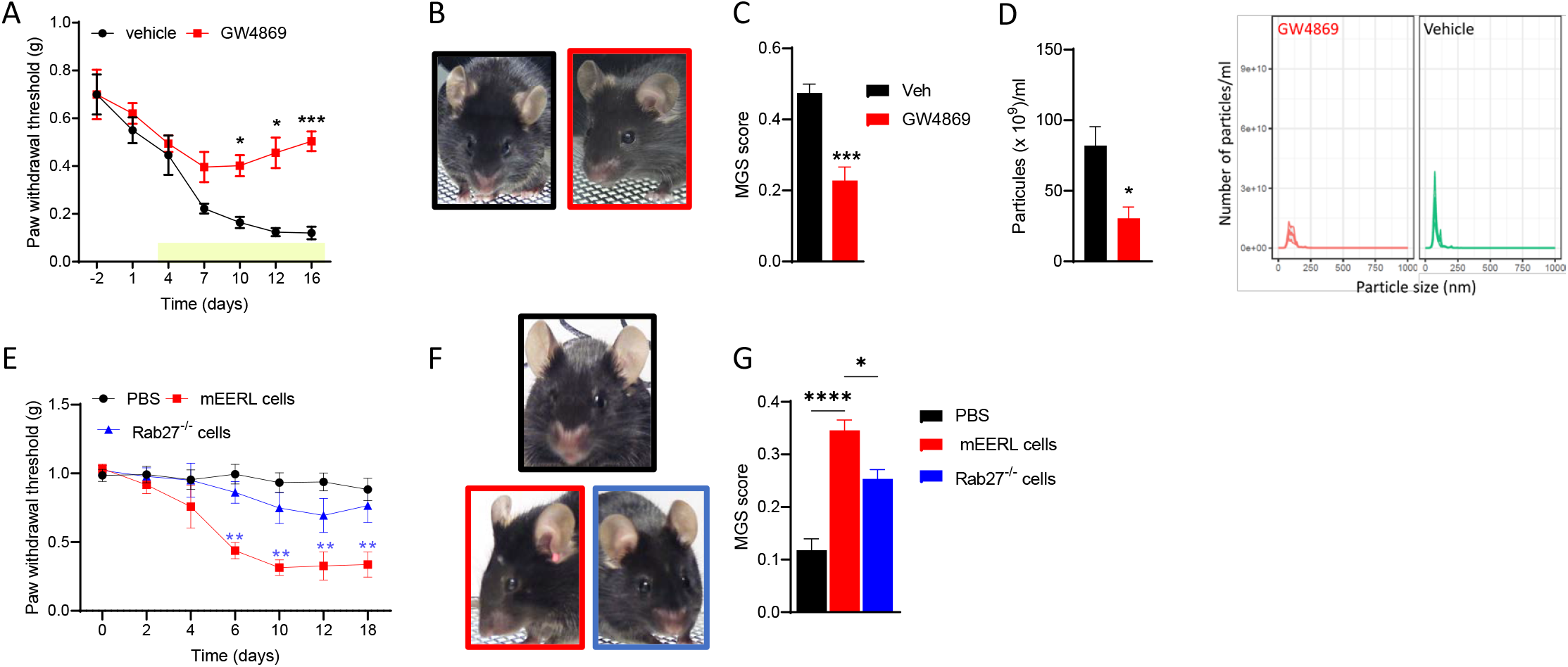
Inhibition of sEV release alleviates cancer pain. (A) GW4869 treatment (beginning on day 3 post tumor implantation, yellow bar) alleviated pain hypersensitivity in tumor-bearing mice (n=5/group; 2-way ANOVA drug effect F (1,8) = 20.2, p<0.002). (B) Representative images of MGS test. (C) GW4869 reduced MGS on day 15 (n=5/group; unpaired t-test 2-tail t=5.4, df=8, p=0.0006) (D) GW4869 reduced the number of circulating particles (n=4/group; unpaired t-test 2- tail t=3.3, df=6, p=0.0162) (E) Genetic deletion of Rab27a and Rab27b in mEERL cells attenuated pain hypersensitivity (n=7/group; 2-way ANOVA cell effect F (2,18) = 47.2, p<0.0001). (F) Representative images of MGS test. (G) Genetic deletion of Rab27a and Rab27b in mEERL cells attenuated MGS (n=7/group; 1-way ANOVA F (2,17) = 32.9, p<0.0001).

### Cancer-derived sEVs are sufficient to induce pain hypersensitivity

Since compromising the release of mEERL-derived sEVs attenuates cancer pain, next we determined whether injection of purified sEVs is sufficient to produce pain hypersensitivity in WT mice. The size and quality of purified sEVs were assessed by nanosight and western blot (**Figure 2A,B**). First, to confirm that isolated sEV injected into the hind paw are taken up by DRG neurons, sEVs were stained blue prior to injection. DRGs harvested 1, 3 and 5 hrs post-injection showed an uptake of the blue dye confirming that sEVs and/or their contents trafficked from the paw to the DRG (**Figure 2C**). Intraplantar injections of sEVs also induced pain hypersensitivity up to 24 hours in a dose-dependent manner (**Figure 3A**). Injection of isolated sEVs have similar effects in male and female mice (**Figure 3B**). Pain hypersensitivity in response to injection of isolated sEVs is not reduce by injection of ketoprofen (clinically used non-steroid anti-inflammatory drug) at a dose classically used in preclinical models (Bagdas et al. 2016) (**Figure 3C**). In contrast, co-treatment with QX-314, an TRPV1+ neuron inhibitor (Binshtok et al. 2007, Talbot et al. 2015), alleviates pain hypersensitivity induced by injection of sEVs into the paw (**Figure 3D**).

**Figure 2.**
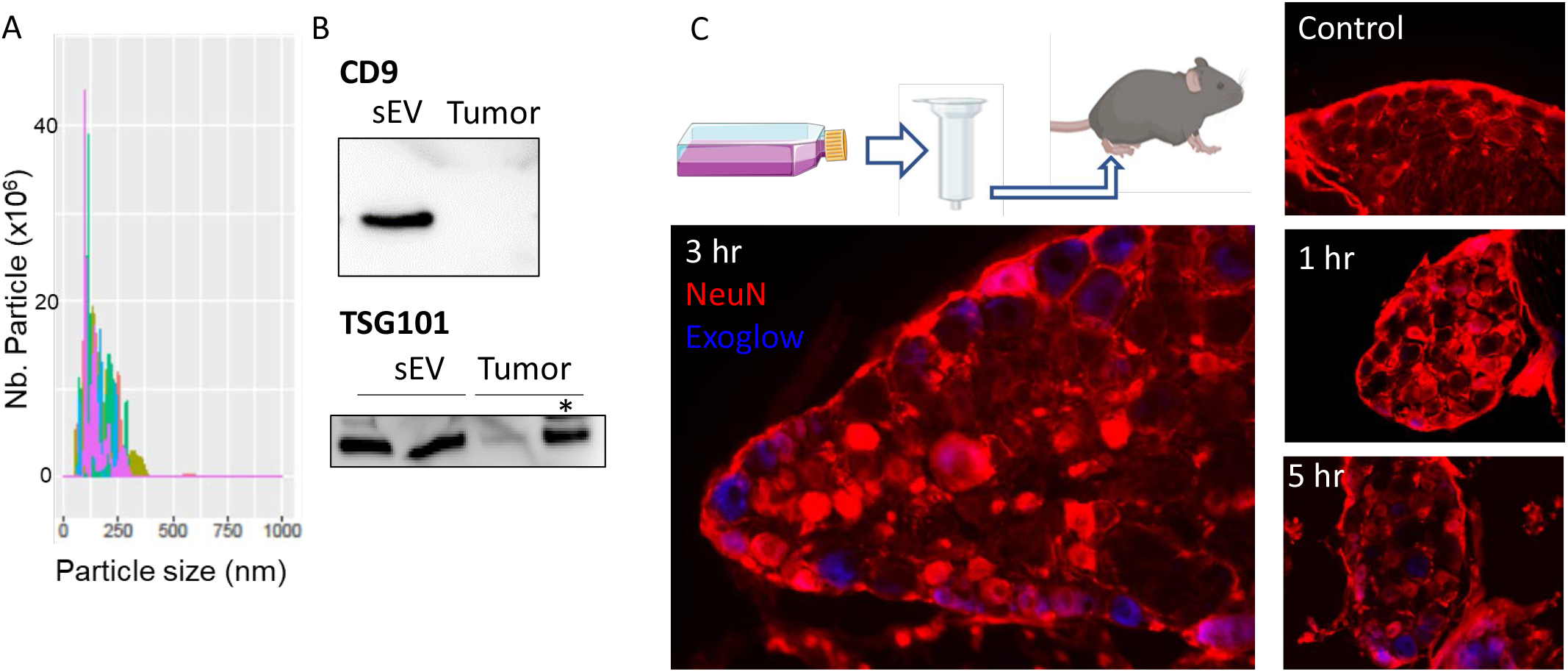
Isolated mEERL-derived sEVs reached the sensory neurons. (A) Representative histogram plot of purified sEVs.by the Nanosight. (B) Representative image of the protein level of CD9 and TSG101 in protein extract from purified sEVs and tumor. Two μg of protein, except * = 30 μg. (C) Diagram of sEV isolation and intraplantar injection. Lumbar dorsal root ganglion sections of PBS- (Ctrl) and Exoglow stained sEV-injected mice.

**Figure 3.**
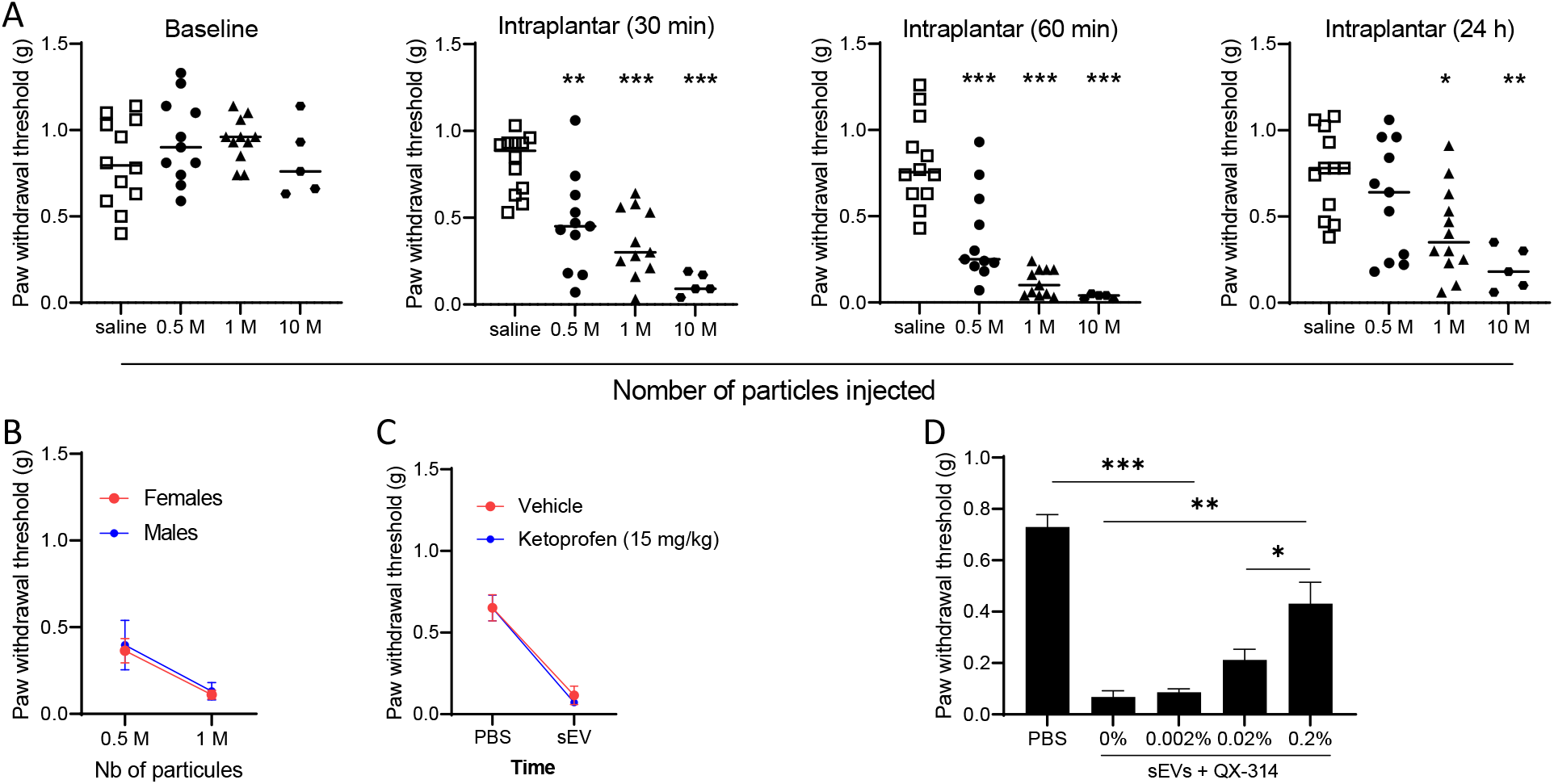
Isolated mEERL-derived sEVs induced pain hypersensitivity in naïve mice. (A) Mechanical sensitivity monitored after intraplantar injection of sEVs (0; 5×10^5^; 10^6^, and 10^7^ particles). Data are compared to saline. One-way ANOVA 30 min F (3,35) = 16.6, p<0.0001; 60 min F (3,35) = 27. 98, p<0.0001; 24h F (3,36) = 6.4, p=0.0013. (B) No sex difference in mechanical sensitivity induced by mEERL-derived sEVs 60 min after injection (n=5-7/group). (C) No analgesic effect of ketoprofen in sEV-injected mice (1M particles, 1 h, n=7/group). (D) QX-314 relieved pain hypersensitivity induced by injection of mEERL-derived sEVs (10^6^ particles, 1 h) (n=8/group, One-way ANOVA F (4,35) = 32.9, p<0.0001.

### Cancer-derived sEVs induce ATF3 expression in TG neurons

To test whether cancer-derived sEVs directly alter sensory neurons, we measured the expression of activating transcription factor 3 (ATF3) a stress-induced protein and neuronal injury marker associated with neuropathic pain (Tsujino et al. 2000, Obata et al. 2003, Peters et al. 2005). First to confirm that cancer cells communicate to sensory neurons by soluble factors, cultured TG neurons are exposed to conditioned media from cultured mEERL cells or control media. Cultured TG neurons showed an induction in ATF3 when exposed to mEERL cell conditioned media for 24 h (**Figure 4A**). Second, TG neurons from male and female mice are exposed to purified sEVs (20 μL equivalent to 3.6 μg of protein) for 24 h. Purified sEVs upregulated ATF3 similarly in female and male TG neurons (**Figure 4B,C**). ATF3 was also upregulated *in vivo* in the sensory neurons of mice implanted with mEERL cells as well as mice that received intraplanar injections of isolated sEVs (**Figure 4D**).

**Figure 4.**
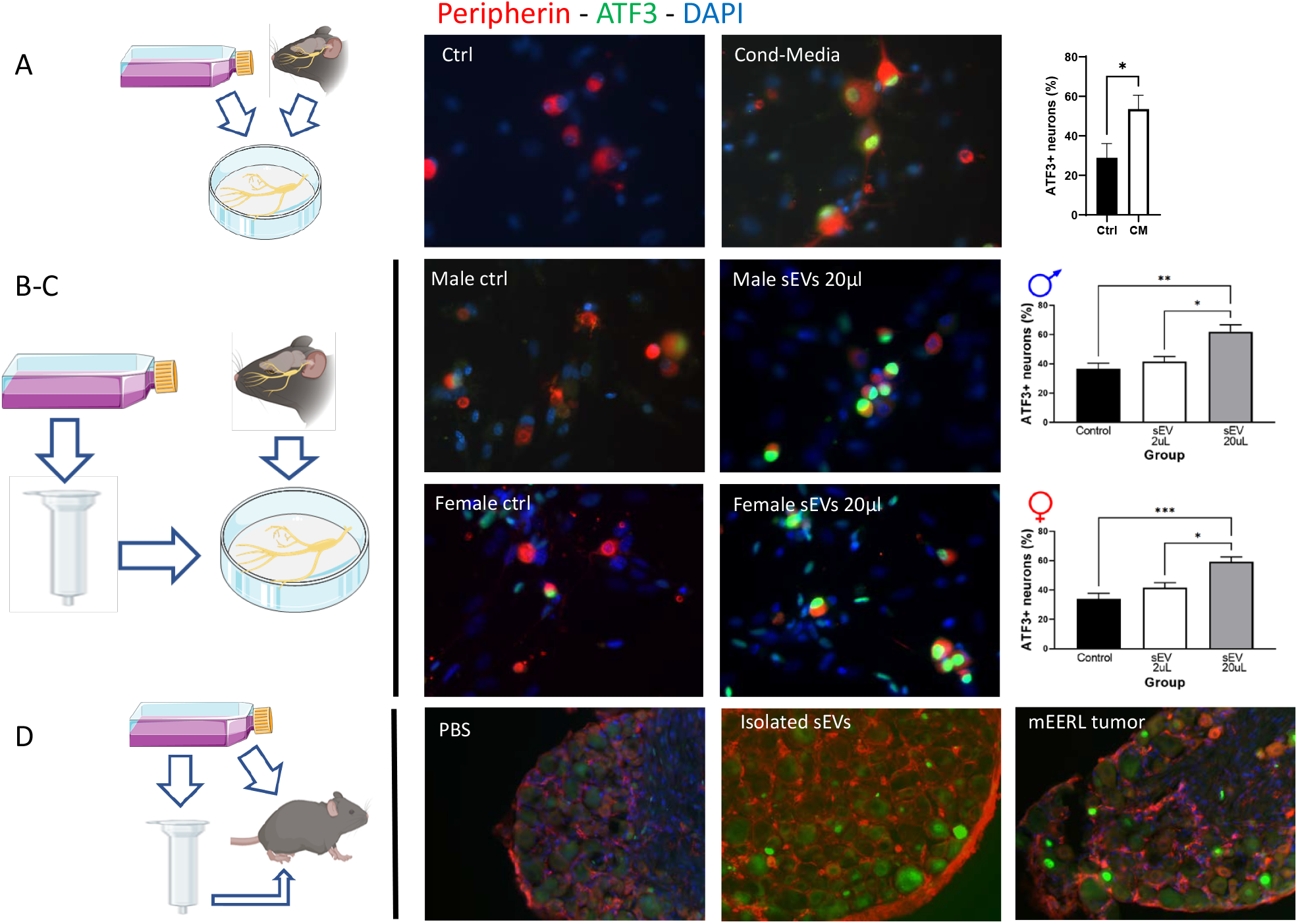
mEERL-derived sEVs induce ATF3 expression in sensory neurons. (A) Cultured TG neurons stained for ATF3 following 24 h exposure to mEERL cell conditioned media. (B) Cultured TG neurons from male mice stained for ATF3 following 24 h exposure to purified sEVs from mEERL cells. (n=5-9/ group, One-way ANOVA F (2,18) = 10.06, p=0.0012 (C) Cultured TG neurons from female mice stained for ATF3 following exposure to purified sEVs from mEERL cells. (n=5-9 neurons/ group, One-way ANOVA F (2,18) = 14.73, p= 0.0002 (D) ATF3 increased in DRG following intraplantar injections of sEVs as well as implantationof mEERL cells.

### Cancer-derived sEVs sensitize TRPV1 neurons

Because sEVs affect cultured TG neurons and induce pain hypersensitivity that is blocked by TRPV1+ neuron inhibitor, we assessed the impact of isolated sEVs by calcium imaging on cultured TG neurons from *Trpv1* ^Cre:^ GCaMP6 mice. Cultured TG neuron showed spontaneous calcium influx as indicated by “fluorescence flashing”. Peaks indicating calcium influx are more frequent and larger in mEERL-derived-sEV-treated cultures indicating that cultured TG neuron incubated with cancer-derived sEVs showed enhanced intracellular calcium events following 24-hr sEVs treatment compared to control TG neurons treated with PBS (**Figure 5 and supplemental videos**). This suggests that sEVs cause TRPV1+ neuron sensitization.

**Figure 5.**
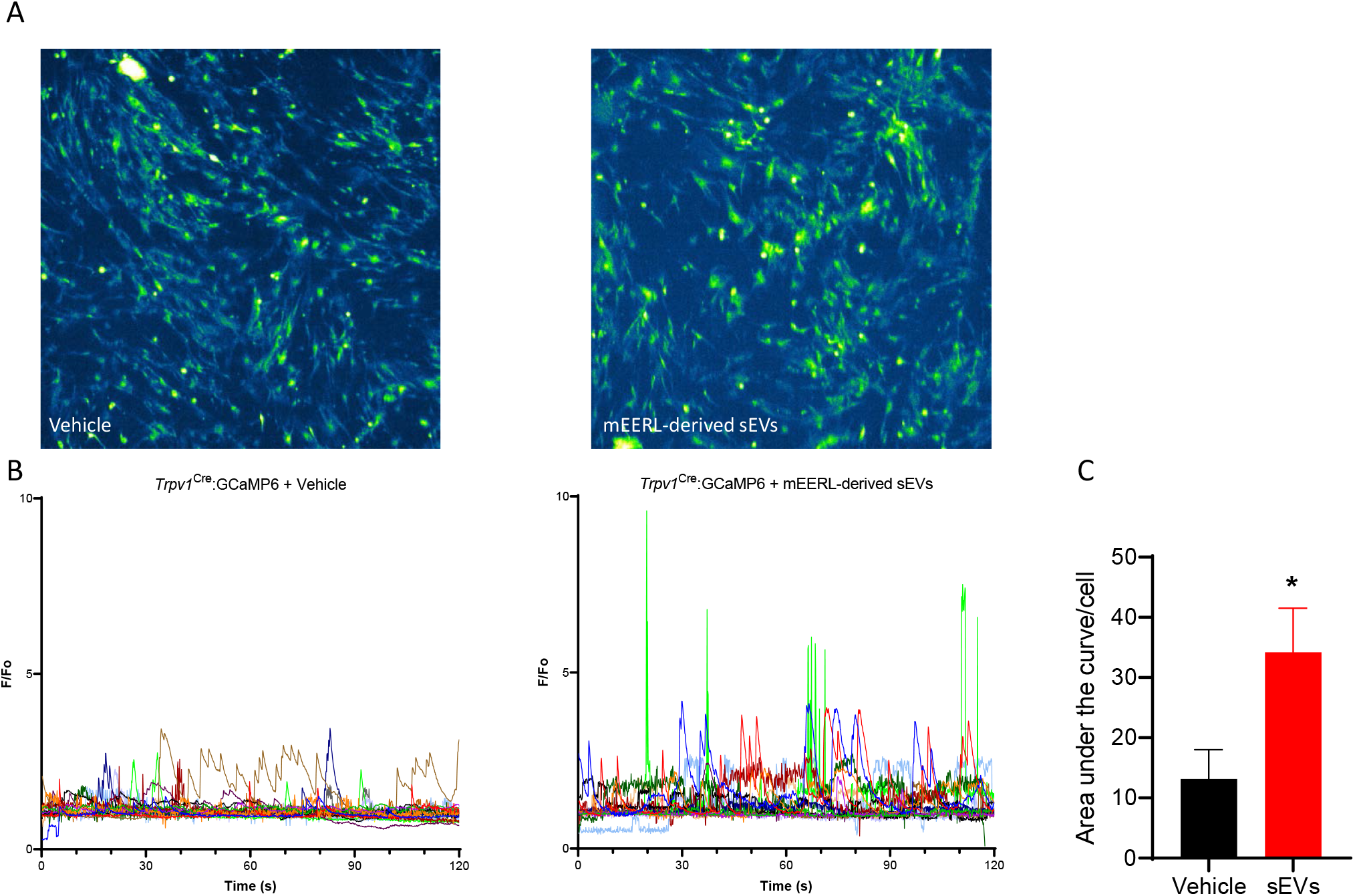
Cultured TRPV1 neurons sensitized by cancer-derived sEVs. (A) Representative images of cultured TG from *Trpv1* ^Cre:^ GCaMP6. (B)Cancer-derived sEVs induced an increase in cellular calcium events in TRPV1+ neurons. Each color line represents a different TRPV1+ cells. (C) Quantification of calcium events: area under the curve of graphs in (B) (n=14-16 cells/groups, Mann-Whitney p=0.013).

### Ablation of TRPV1+ neurons prevent cancer pain

In order to assess the role of TRPV1+ neurons on cancer pain, naïve mice were treated with RTX to ablate that TRPV1-expressing neurons. The ablation was confirmed by the absence of reaction to the hot plate test (**Figure 6A**) and a lack of TRPV1+ neurons in sensory ganglia (**Figure 6B,C**). Four weeks after RTX treatment, mice were injected with mEERL cells into the right hindleg. Von Frey (**Figure 6D**) and mouse grimace scale testing (**Figure 6E,F**) indicated that RTX treatment prevents evoked and spontaneous cancer pain but did not affect baseline mechanical sensitivity.

**Figure 6.**
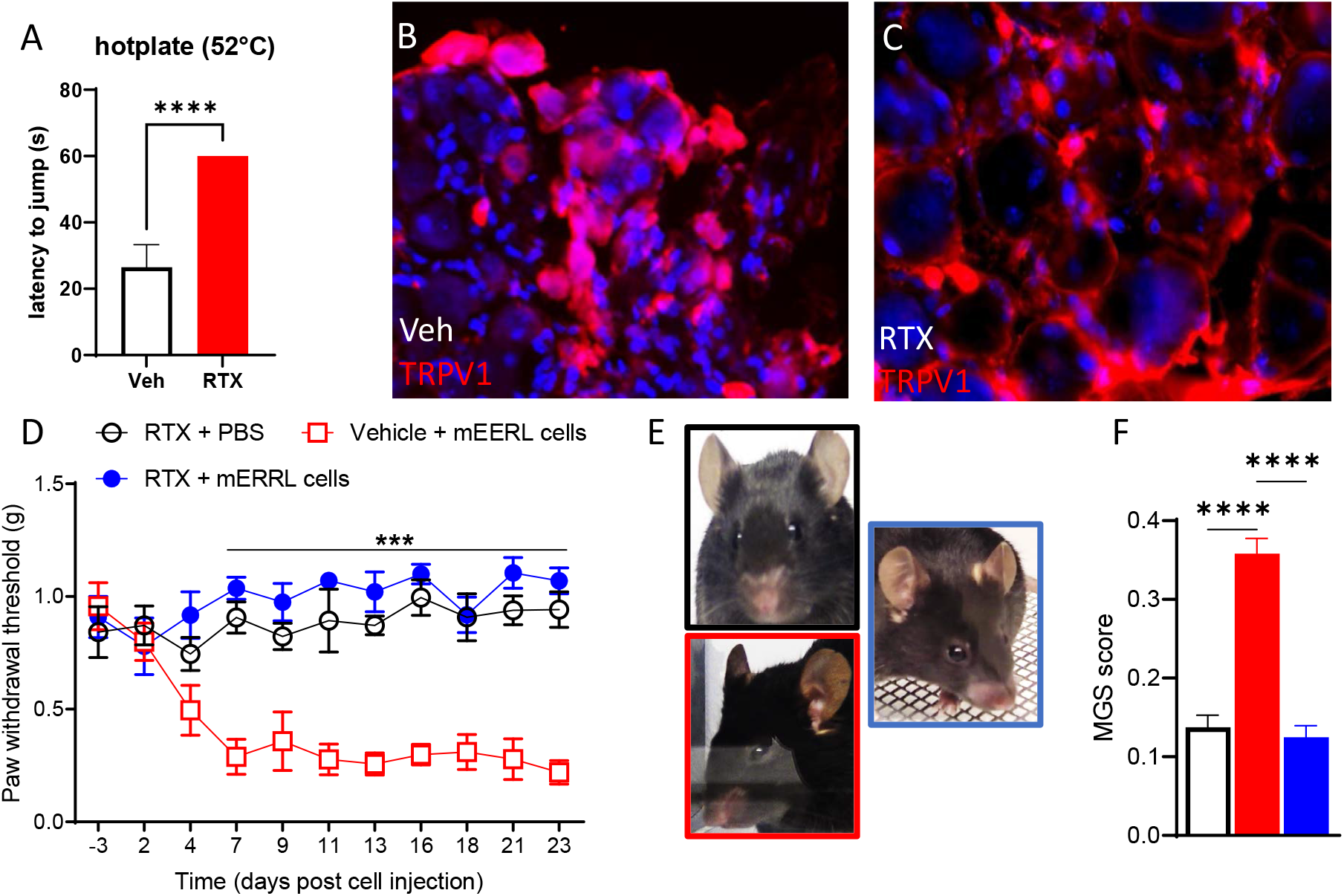
Chemo-ablation of TRPV+ neurons prevents cancer pain. (A) RTX-treated mice are unresponsive to the hotplate (veh n=6, RTX n= 13; unpaired t-test t=18.3, df=17, P<0.0001). (B-C) Representative images of TRPV1 staining in the dorsal root ganglion. (D) Chemo-ablation of TRPV1+ neurons blocks pain hypersensitivity in tumor-bearing mice (n=6-8/group, 2-way ANOVA treatment effect F (2,17) = 112.5, p<0.0001). (E) Representative image of mouse facial grimacing. (F) Chemo-ablation of TRPV1+ neurons alleviates facial grimacing in tumor-bearing mice (n=6/group, one-way ANOVA F (2,15) = 63.4, p<0.0001).

### Cancer-derived sEVs activate the translation initiation pathways to induce pain hypersensitivity

To further investigate the potential mechanism of nociception induced by HNSCC-derived sEVs and identify clinically relevant targets, we took advantage of publicly available human RNA-sequencing data. RNA sequencing performed from unstimulated cultured human DRGs (5 different cultures) (Wangzhou et al. 2020) and exposed to human HNSCC-derived sEVs (3 cultures) (Amit et al. 2020). After removing genes that were not present in both datasets and had missing expression data, we generated a list of 9369 genes. We selected the genes with 0.2>log2FC>3 and a p-value<0.001 to obtain 1716 genes (**Supplementary table**). To infer the functional role of these differentially expressed genes, we performed gene-enrichment pathway analysis. Ingenuity Pathway analysis (IPA) data indicate that sEVs induce several canonical pathways linked with the initiation of protein translation such eukaryotic initiation factor (eIF) 2 and 4 signaling, mammalian target of rapamycin (mTOR) signaling, and p706K signaling. (**Table 1**). All of these pathways, involved in the regulation of the nascent translation, contribute to nociception (Price et al. 2009, Sonenberg et al. 2009). To test whether cancer-derived sEVs also trigger the translation of nascent protein in our mouse model, we used the methionine analog AHA to label newly translated proteins in cultured TG neurons (Melemedjian et al. 2010). Incubation with sEVs increased AHA fluorescence in peripherin+ cells (TG neurons) indicating an enhancement of translation of nascent proteins (**Figure 7A,B**), aligning our preclinical model with human sequencing data: cancer-derived sEVs induced translation of nascent protein in sensory neurons. These pathways are blocked by AMP-activated protein kinase (AMPK) activation and mTOR inhibition (Hardie 2007, Melemedjian et al. 2013, Inyang et al. 2019). Consistently, in tumor bearing mice, inhibition of mTOR directly through rapamycin attenuates pain hypersensitivity (**Figure 7C**). Moreover, administration of AMPK activator narciclasine (NCLS) (Zhang et al. 2009, Julien et al. 2017) prevents evoked and spontaneous cancer pain (**Figure 7D-F**).

**Figure 7.**
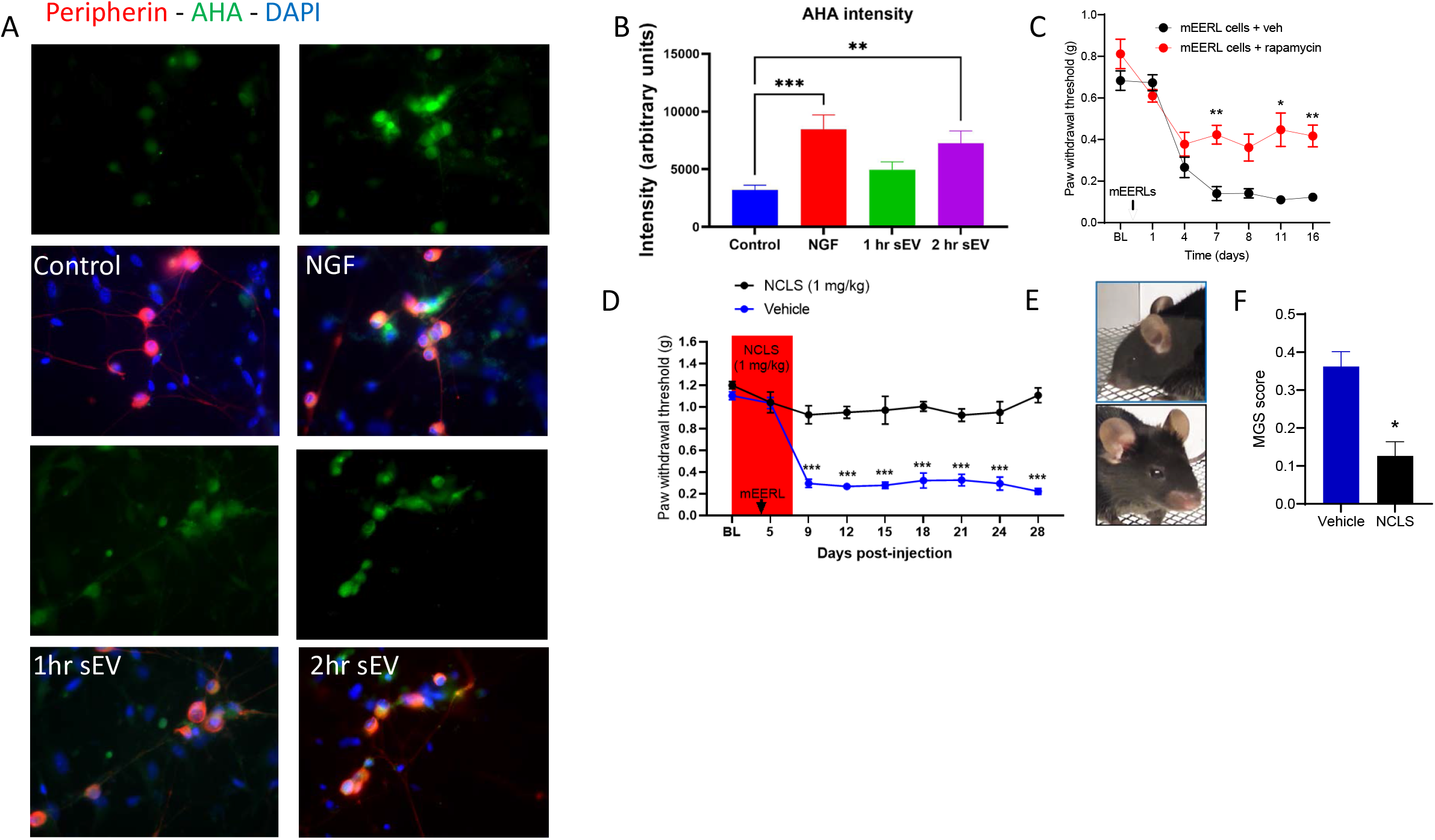
Blocking the translation initiation pathways alleviates cancer pain. (A) Representative images of intensity correlation analysis of fluorescently labeled AHA click iT chemistry with neuronal markers (B) sEVs induce translation of nascent proteins in cultured TG neurons at 2 hrs. (n=20- 23 cells/group, One-way ANOVA F (3,85) = 6.873, 2 hr sEV p = 0.0074). (C) Rapamycin treatment attenuated cancer-induced mechanical hypersensitivity (n=7/group, 2-way ANOVA rapamycin effect F(1,12) = 33.1, p<0.0001). (D) Narciclasine treatment prevents cancer-induced mechanical hypersensitivity (n=4- 5/group, 2-way ANOVA NCLS effect F (1,63) = 385.4, p<0.0001). (E) Representative images of mouse facial grimacing. (F) Narciclasine treatment alleviate facial grimacing in tumor-bearing mice (n=4/group, Mann-Whitney p=0.03).

## Discussion

One of the novel key findings of this study is that TRPV1+ neurons are directly sensitized by cancer-derived sEVs and these sEVs are necessary for cancer pain. Cancer-derived sEVs induced translation of nascent proteins in sensory neurons to mediate cancer pain.

Chemo-ablation of TRPV1+ neurons completely prevents the development of cancer pain indicating a major role of TRPV1+ neurons. Other reports have shown that RTX treatment alleviates cancer pain in preclinical, veterinarian, and clinical studies (Heiss et al. 2015, Sapio et al. 2018). Given the promising data, a phase II clinical trial is ongoing to test whether intrathecal RTX reduces cancer pain (National Institute of et al. 2023). Nonetheless, using genetically specific calcium imaging our study is the first to show that TRPV1+ neurons are directly sensitized by cancer-derived sEVs.

The reason cancer cells communicate with TRPV1+ neurons remains elusive, but is the subject of intense investigation (Demir et al. 2021). This field was opened by the groundbreaking discovery that ablation of TRPV1+ neurons slowed down the progression of pancreatic cancer (Saloman et al. 2016). One reason might be the regulation of tumor immunology by neurons (Scheff et al. 2022, Udit et al. 2022). Given that pain is a prognostic factor for survival (Reyes-Gibby et al. 2014), it suggests shared mechanisms linking carcinogenesis and pain in HNSCC (Ye et al. 2022). Our present work shows that sEVs released by cancer cells are a key mediator of communication to TRPV1+ neurons in line with previous reports (Madeo et al. 2018, Vermeer 2019, Amit et al. 2020). Alternative mechanisms to activate TRPV1+ neurons cannot be disregarded such as lower pH in the tumor environment, nerve growth factor (NGF) and Ephrin produced by cancer cells are well known inducers of nociception (Ye et al. 2011, Madeo et al. 2018) and direct activation of TRPV1+ neurons (Scheff et al. 2022). However, pharmacological, and genetic attenuation of cancer-derived sEV release alleviate cancer pain in tumor-bearing mice. This result coupled with the fact that injection of isolated sEVs trigger pain hypersensitivity indicates that sEVs play a critical role in cancer pain. Previous publication reports that injection of human isolated cancer exosomes induced pain hypersensitivity in mice (Bhattacharya et al. 2020), but the potential immune response to human antigen could not be excluded. Here, we showed that sEVs from cancer cells with a C57 background induces pain hypersensitivity in mice from the same genetic background. Additionally, our data indicate that injection of the anti-inflammatory pain killer ketoprofen does not attenuates pain induced by sEV injection demonstrating that potential inflammation in response to sEVs is not driving the pain and that sEVs are not likely to induce inflammation.

While cancer-derived sEVs are a key player in regulation of cancer pain, it remains unclear how sEVs sensitize neurons. sEVs can release their cargo in neurons by membrane fusion or endocytosis to affect neuronal physiology. Additionally, sEV transmembrane proteins or lipids may bind to receptors expressed on TRPV1+ neurons. We discuss several mechanisms potentially at stake.

The ability for HNSCC-derived sEVs to induce axonogenesis is well established (Lucido et al. 2019, Amit et al. 2020). Axon growth is likely to be associated with major physiological changes including transcriptional and translational. Upregulation of ATF3 in neurons following exposure to isolated sEVs further supports this hypothesis. ATF3 is classically upregulated after nerve injury, TRPV1 activation, and is associated with transcriptional reprogramming necessary for peripheral nerve regeneration and neuropathic pain (Tsujino et al. 2000, Bráz et al. 2010, Renthal et al. 2020), suggesting that cancer pain has a neuropathic component (Ye et al. 2022). The upregulation of ATF3 in response to sEVs supports the idea that sEVs drastically impact the neuronal gene expression to switch to a transcriptome facilitating axon elongation and neuronal sensitization. Additionally, sEVs are known to be filled with miRNAs (Akers et al. 2013, Tkach et al. 2016, Mao et al. 2018). These miRNAs may affect neuronal epigenome, gene expression, and translation and contribute nociception. Epigenetic changes and miRNAs contribute to the chronification of pain (Sakai et al. 2013, Laumet et al. 2015, Pan et al. 2016, Peng et al. 2017). The exact contribution of sEV-derived miRNAs to cancer pain is unclear and will be further explored. Moreover, observation from human RNA-seq data and our in vitro model point out that that sEVs induce eIF pathway/nascent translation signify that in addition to transcriptional reprogramming, cancer-derived sEVs trigger initiation of translation in sensory neurons. ATF3 also plays a key role in axonal translation (Jiang et al. 2004). Nociception resulting from enhanced nascent translation is observed across pain models and supported by human data (Melemedjian et al. 2010, Khoutorsky et al. 2018) indicating that targeting translation inhibition is promising for cancer pain.

Activation of the eIF pathway /nascent translation may result from activation of Protease activated receptor 2 (PAR_2_), a G-protein-coupled receptor expressed by nociceptors, that mediates pain (Steinhoff et al. 2000, Vergnolle et al. 2001) in an eIF4F-dependant fashion (Tillu et al. 2015), and its activation results in an increase in ATF3 expression (Falconer et al. 2019). Then PAR_2_ sensitizes and activate TRPV1 (Amadesi et al. 2006). Tissue factor (TF), a PAR_2_ ligand, may be released by extracellular vesicles (Gardiner et al. 2015, Date et al. 2017). Studies showed that PAR_2_ plays a critical role in HPV- oral cancer pain (Lam et al. 2012, Tu et al. 2021). PAR_2_ and TRPV1 might be also activated by lipid (Lam et al. 2012, Ruparel et al. 2015, Tu et al. 2021) released by sEV or included in their membranes. Alternatively, tumor-secreted NGF may also trigger the eIF pathway /nascent translation (Melemedjian et al. 2010). Activation of mTOR pathways may also facilitate tumor innervation (Madeo et al. 2018, Vermeer 2019, Wong et al. 2022).

Interesting despite the strong sexual dimorphism in chronic pain prevalence (Mogil 2012), pain in response to the sEVs is similar in both sexes. Consistently, ATF3 upregulation is also similar in TG cultured neurons from female and male mice. One important limitation is that sEVs are isolated from mEERL cells that have been isolated from male mice. If no sex difference is observed with male sEVs affecting equally female and male neurons, a difference may emerge between sEV isolated from female and male mice. However, the presence of sexual dimorphism in HNCP is still elusive (Ye et al. 2022).

Our study provides several promising potential targets for treating HNSCC pain. In addition to RTX that is already in clinical trials (National Institute of et al. 2023), QX-314, a modified lidocaine specific to TRPV1+ neurons, is a promising target for cancer pain. Interfering with sEV release from cancer cells is a valuable therapeutic strategy. Activating AMPK to block the EIF pathways/nascent translation might be the most promising because AMPK activators like NCLS inhibits metastasis and tumor growth as well (De Benedetti et al. 2004, Yousuf et al. 2021) and AMPK activators are well known for their analgesic effects and some like metformin are already FDA-approved (Khoutorsky et al. 2018).

In summary our work reveals that HNSCC-derived sEVs sensitize TRPV1+ neurons by promoting nascent translation to mediate cancer pain and identified several promising therapeutic targets to interfere with this pathway.

### STAR Method

All animal experiments were approved by MSU IACUC and in accordance with NIH guidelines.

|  | Source | Identifier |
| --- | --- | --- |
| **Animal** |  |  |
| C57Bl/6J mice | Jackson laboratories | JAX# 000664 |
| GCaMP6 | Jackson laboratories | JAX# 031968 |
| Trpv1<sup>Cre</sup> | Jackson laboratories | JAX# 017769 |
| **Cells** |  |  |
| mEERL | Paola Vermeer | Madeo et al. 2018 |
| mEERL *Rab27a*<sup>-/+</sup> and *Rab27b*<sup>-/-</sup> | Paola Vermeer | Madeo et al. 2018 |
| **Drugs** |  |  |
| Resiniferatoxin | Sigma-Aldrich | R8756 |
| Narciclasine | Santa Cruz | sc-361271 |
| Dimethyl sulfoxide | Fisher Scientific | D128-1 |
| 2-Hydroxypropyl-β-cyclodextrin | Santa Cruz | sc-203461A |
| Rapamycin | Sigma-Aldrich | #37094 |
| GW4869 | Sigma-Aldrich | D1692 |
| QX-314 bromide | Tocris Biosciences | #1014 |
| Ketoprofen | Sigma-Aldrich | K1751 |
| **Antibodies** |  |  |
| Anti-CD9 | Abcam | ab263019 |
| Anti-TSG101 | Abcam | ab125011 |
| Anti-Rabbit | Abcam | ab205718 |
| Anti-ATF3 | Abcam | ab207434 |
| Anti-TRPV1 | Abcam | ab203103 |
| Alexa Fluor anti-rabbit 488 | Thermofisher | A-11008 |
| Alexa Fluor anti-mouse 568 | Thermofisher | A-11004 |
| Anti-peripherin | Sigma-Aldrich | P5117 |
| **Chemicals** |  |  |
| L Azidohomoalanine (AHA) | Invitrogen | C10102 |
| Nerve growth factor | Sigma-Aldrich | #93928-24-6 |
| **Enzymes** |  |  |
| collagenase A | Sigma-Aldrich | #10103578001 |
| collagenase D | Sigma-Aldrich | #11088858001 |
| trypsin inhibitor | Sigma-Aldrich | #10109886001 |
| **Commercial assays** |  |  |
| ExoGlow-Protein EV Labeling Kit | System Biosciences | EXOGP400A-1 |
| ExoQuick-TC | System Biosciences | EXOTC10A-1 |
| VIVASPIN 100kDa ultrafiltration tubes | Sartorius | VS2001/VS2041 |
| IZON columns | Izon | SP2 |
| Amicon 10kDa centrifugal filter tube | Millipore Sigma | UFC801024 |
| **RNA-seq data** |  |  |
| Human cultured DRG | https://www.ncbi.nlm.nih.gov/pmc/articles/PMC7305999/ | Wangzhou et al. 2020 |
| Human cultured DRG + sEVs | https://www.ncbi.nlm.nih.gov/geo/query/acc.cgi?acc=GSE134220 | Amit et al. 2020 |
| Ingenuity Pathway Analysis | QIAGEN |  |
| **Calcium imaging** |  |  |
| Andor Zyla 4.2 PLUS scMOS digital camera | Oxford Instruments |  |
| stereomicroscope | Nikon | SMZ18 |
| Cell Profiler 4.0 | Stirling et al., 2021 |  |

The full description of the materials and methods is available on the **Supplementary information**.

## Acknowledgement

This work was supported by the Rita Allen Foundation (G.L.), the NIH NINDS 1R0121259 (G.L.)

We thank Greg Dussor and Theodore Price (The University of Texas at Dallas) for expert advice on trigeminal ganglion dissection and click chemistry respectively, Issac M. Chiu and Daping Yang (Harvard University) for guidance for RTX-neuronal ablation, Cole McCutcheon (MSU) for help and support with the Nanosight, and Karli Monahan (MSU) for technical assistance.

## Inclusion and Diversity

We worked to ensure sex balance in the selection of non-human subjects. One or more of the authors of this paper self-identifies as an underrepresented ethnic minority in science. One or more of the authors of this paper self-identifies as a member of the LGBTQ+ community. While citing references scientifically relevant for this work, we also actively worked to promote gender balance in our reference list.

## Author contributions

KEI, CME, MH, JKF, and GL performed experiments. MP guided sEV isolation. MR performed bioinformatics. NT performed calcium imaging experiments. KEI, PDV, NT, JKF, and GL analyzed data. KEI, JKF, PDV, and GL conceived the study. KEI, JKF, and GL wrote the manuscript. GL oversaw the study. All authors approved the final version of the manuscript.

The authors declare no competing interests

## Figure legends

**Supplementary figure 1.**
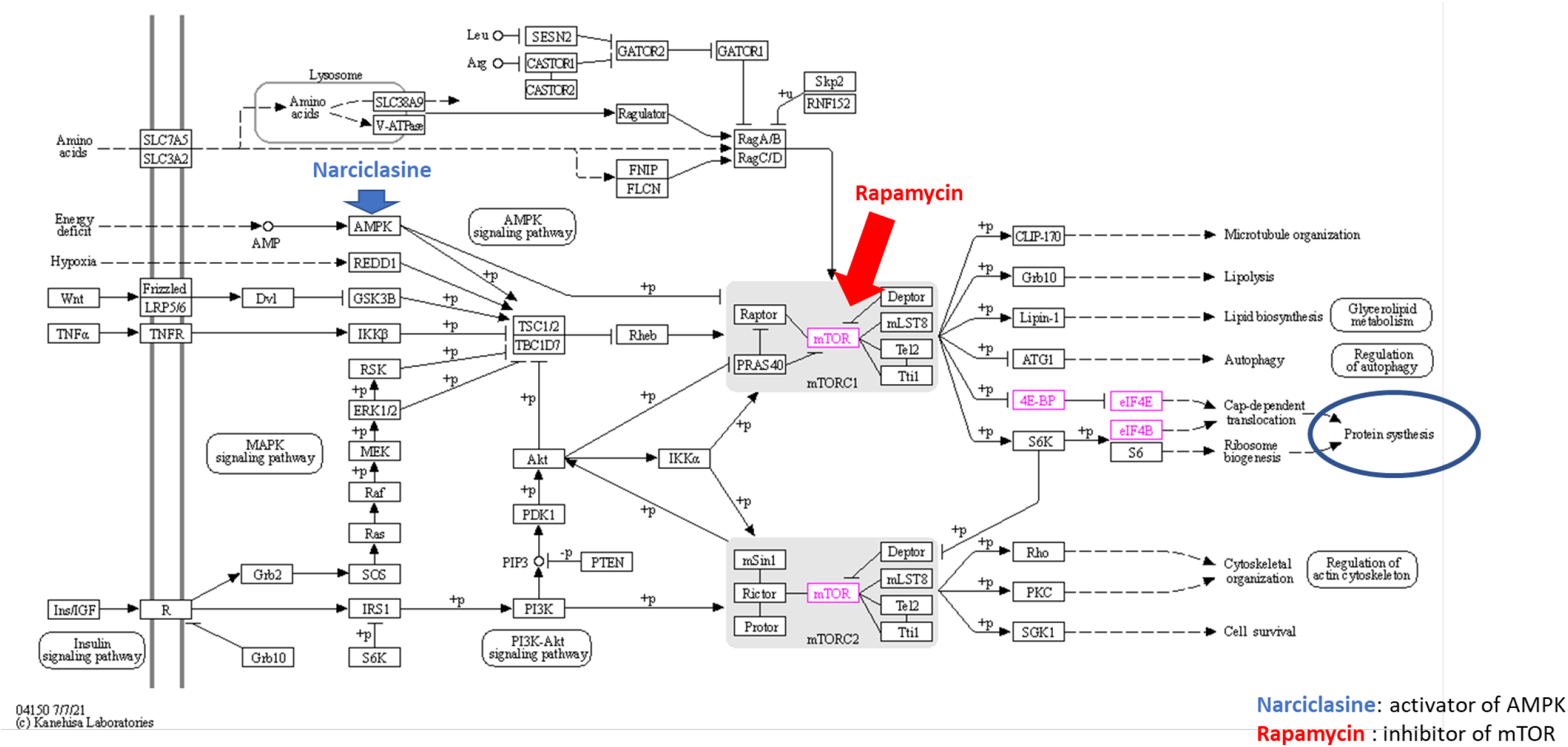
AMPK/mTOR/eIF signaling pathways promoting protein translation. Rapamycin is an mTOR inhibitor and narciclasine is an AMPK activator.

