## Supplemental methods for "Head and Neck Cancer-derived small extracellular vesicles sensitize TRPV1+ neurons to mediate cancer pain"

Geoffroy Laumet

Department of Physiology

College of Natural Science

Michigan State University

Interdisciplinary Science and Technology Building

766 Service Rd

East Lansing, MI 48826, USA

Keywords: cancer, head and neck squamous cell carcinoma (HNSCC), pain, extracellular vesicles, TRPV1, translation, AMPK

**Experimental procedures**

Animal procedures were approved by Michigan State University Institutional Animal Care and Use Committee (IACUC) and were in accordance with National Institutes of Health Guidelines. Experiments were performed on male and female C57Bl/6J mice (JAX# 000664, Jackson Laboratory, Bar Harbor, ME) except those related to calcium imaging. For the latter, *B6.129-Trpv1^tm1(cre)Bbm^/J* (JAX# 017769) (Cavanaugh et al. 2011) were bred with *6(129S4)-Gt(ROSA)26Sor^tm1.1(CAG-tdTomato/GCaMP6f)Mdcah^/J* (JAX# 031968) (Dong et al. 2017) mice to obtain genetically encoded calcium indicator GCaMP6f-tdTomato fusion protein expression specifically in TRPV1+ neurons (*TRPV1^Cre^:Salsa6f* mice). These mice were subsequently called *TrpV1*^Cre^:GCaMP6. Mice were bred and housed in the Michigan State University Animal Care Facility prior to the start of behavioral testing. Animals had *ad libitum* access to food and water and were on a 12 hr non-inverted light/dark cycle. Mice were randomized to treatment groups.

HNSCC model: To produce HNSCC tumors, we used a well-established murine model of human papillomavirus induced (HPV +) oropharyngeal squamous cell carcinoma (Hoover et al. 2007, Spanos et al. 2008, Madeo et al. 2018, Heussner et al. 2021). This model consists of oropharyngeal epithelial cells from C57Bl/6 male mice that stably express HPV16 viral oncogenes, E6 and E7, activated H-Ras and luciferase (mEERL cells). These cells were cultured and injected subcutaneously into the right hindleg of 10-16-week-old male mice to produce tumors. Injection into the leg instead of the oral cavity is justified by the 3Rs guideline in an effort to reduce distress, inability to feed, and easier assessment of tumor growth. Only male mice were injected with tumor cells because the cell line was generated from male mice and cannot be implanted in immunocompetent female mice.

Generation of mEERL *Rab27a*^-/+^ and *Rab27b*^-/-^ cells: to compromise the release of sEVs in cancer cells, *Rab27a* and *Rab27b*, two genes encoding proteins essential for exosome release, were knock-out using CRISPR-Cas9 technology in mEERL cells. The gene editing approach and attenuation of sEV release were validated in a previous publication (Madeo et al. 2018).

mEERL culture: mEERL cells were cultured in Dulbecco’s modified Eagle’s medium F-12 GlutaMax media (#10565018; Fisher Scientific, Hampton, NH) that contained 10% fetal bovine serum (FBS; #26-140-079; Fisher) and 1X penicillin streptomycin (Pen-Strep; #15070063; Fisher) as previously described (Heussner et al. 2021).

Behavioral testing: was performed by experimenters blinded to experimental conditions. Mechanical hypersensitivity: was assessed by stimulation of the hindpaw with calibrated von Frey filaments from Stoelting using the Dixon up-down method (Dixon 1965) following 45 min of habituation to the testing boxes as previously described (Heussner et al. 2021, Inyang et al. 2021). Mouse grimace scale (MGS): assesses non-evoked pain using facial features (Langford et al. 2010) such as orbital tightening, cheek bulge, nose bulge, and ear position. Each feature is evaluated on a 3-point scale, and then the sum is analyzed as previously described (Heussner et al. 2021). Hotplate: Mice were individually habituated in the apparatus with the plate set to 30^o^C for 2 min. The temperature of the hotplate was then increased to 52^o^C. Mouse was placed on the plate and the latency to jump within 60 sec was recorded. If mice did not jump, the score was 60 sec.

TRPV1+ neuron ablation**:** Subcutaneous injection of resiniferatoxin (RTX, R8756; Sigma- Aldrich, St. Louis, MO) in the flank was performed on 4-week-old mice with escalating doses of 30 μg/kg at day 1, 70 μg/kg at day 2 and 100 μg/kg at day 3. Control mice received a vehicle solution consisting of DMSO with Tween80 in phosphate-buffered saline (PBS). Denervation was evaluated using the hot plate method (Karai et al. 2004, Pinho-Ribeiro et al. 2018) and immunostaining for TRPV1.

Small extracellular vesicle purification: sEVs were purified following a method previously validated (Nguyen et al. 2021). Briefly, sEVs were harvested from cultured mEERL cells cultured 24 h in exosome-depleted serum (A2720803, Invitrogen, Carlsbad, CA). Culture media was collected, centrifuged and steriflip filtered sterilized. This solution was then placed in VIVASPIN 100kDa ultrafiltration tubes (VS2001/VS2041; Sartorius, Göttingen, Germany) and centrifuge for 50 min at >3000G to concentrate volume down to ~50-150ul. The concentrated solution was run through IZON columns (SP2; IZON Science, Christchurch, New Zealand) and 200 µl fractions were collected. Fractions 6, 7, 8 are enriched for vesicles and have minimal protein contamination. The sEVs were concentrated by adding them to a Amicon 10kDa centrifugal filter tube (UFC801024; Millipore Sigma, St. Louis, MO) and spinning 50 min at >3000G at room temperature. Purified sEVs were quantified using NanoSight NS300 (Malvern Panalytical, Westborough, MA) as described previously (Nguyen et al. 2019). Additionally, the expression of sEV markers is assessed by western blot (**Figure 2**). Isolated sEVs yield ~10^6^ particles/µl or 0.1-0.2 µg of protein/µl).

Western blotting: All supplies from Invitrogen, Carlsbad, CA unless otherwise mentioned. Two ug of total protein from sEV samples were mixed with NuPAGE LDS Sample Buffer (NP0007; ThermoFisher, Waltham, MA) under reducing conditions and loaded onto a NuPAGE Novex Bis-Tris 4-12% pre-cast Gel (NP0321BOX; ThermoFisher). An equivalent sample of protein from a mEERL cell tumor, lysed in RIPA buffer, was also run as a control. Proteins were separated under reducing conditions in NuPAGE MES running buffer for 35 min at 200 V constant voltage. After separation the proteins were then transferred to PVDF membrane using NuPAGE transfer buffer with 10% methanol at 30 V constant voltage for 1 hour. After transfer the membrane was blocked for 1 hour in Pierce Clear milk blocking solution and then incubated at 4^o^C overnight with primary antibody Anti-CD9 antibody [EPR23105-125] (1:2000; ab263019; Abcam, Cambridge, United Kingdom) and Anti-TSG101 antibody [EPR7130(B)] (1:2000; ab125011; Abcam), which was prepared in the Pierce clear milk blocking solution.

The next day the membrane was washed 3 times for 10 min with Tris Buffered Saline with 0.05% Tween20 (TBST) and then incubated for 1 hour at room temperature with a Goat anti-Rabbit IgG antibody with an HRP tag (1:4000; ab205718; Abcam) in TBST. The membrane was then washed 4 times with TBST for 5 min and then once with Tris Buffered Saline (TBS) for 5 min. The membranes were then incubated for 5 min at room temperature with Supersignal West Dura HRP substrate, excess substrate was then drained, and the membrane placed in a clear plastic folder. The membrane was then imaged using a Aplegen Omega Lum G (#8418-10-0005; GMI Inc, Ramsey, MN).

Primary neuron culture: Trigeminal ganglia (TG) were extracted aseptically from 8-week-old male and female mice in Hanks’ buffered salt solution (HBSS; #24020117; Fisher) on ice to be cultured using a previously used procedure (Inyang et al. 2019). The TGs were dissociated enzymatically at 37°C, first with collagenase A (1 mg/ml; #10103578001; Sigma-Aldrich) for 25 minutes and then collagenase D (1 mg/ml; #11088858001; Sigma-Aldrich) that included papain (30 μg/ml; #10108014001; Sigma-Aldrich) for 20 minutes. Afterward, a trypsin inhibitor (1 mg/ml; #10109886001; Sigma-Aldrich) that contained bovine serum albumin (bovine serum albumin, 1 mg/ml; Fisher) was applied, and the ganglia were mixed to allow further dissociation with a polished Pasteur pipette. The tissue was then filtered through 70-μm nylon cell strainer (CLS431751; Sigma-Aldrich) and resuspended in Dulbecco’s modified Eagle’s medium F-12 GlutaMax media (#10-565-018; Fisher Scientific) that contained 10% fetal bovine serum (#26-140-079; Fisher Scientific) and 1X penicillin streptomycin (#15070063; Thermo Scientific, Waltham, MA). The media also contained nerve growth factor (1:1000; #93928-24-6; Sigma-Aldrich). Neurons were cultured for 4 days on Poly-L-Lysine German Glass Coverslips #1, 12 mm (#72292-02; Fisher Scientific) in a 24-well tissue culture plate (#09-761-146; Fisher Scientific) at 37°C with 95% air and 5% CO_2_. On the day of the experiment, purified sEVs were diluted into Dulbecco’s modified Eagle’s medium F-12 plus GlutaMax media and added directly onto the neurons and incubated for 24 hours.

Immunocytochemistry and Digital Image Analysis: After sEVs treatment, the cells were washed with PBS and fixed with 10% formalin in PBS for 30 minutes. Cells were blocked with 10% normal goat serum and labeled with anti-peripherin, mouse monoclonal (1:500; P5117; Sigma) and activating transcription factor 3 (ATF3) (1:1000; ab207434; Abcam) overnight at 4°C. Next, cells were washed and incubated with flurochrome-conjugated secondary antibodies (Alexa Fluor, anti-rabbit 488 (1:1000; A-11008) and anti-mouse 568 (1:1000; A-11004; Thermofisher, Waltham, MA) and counterstained with a DNA stain, 4′,6-diamidino-2-phenylindole (DAPI) (1:50,000; D1306; Fisher) and mounted with Prolong Gold (P36930; Invitrogen, Carlsberg, CA). Staining was visualized using a fluorescent microscope (Nikon Eclipse Ni-U, Minato City, Tokyo, Japan) and images were created using ImageJ (National Institute of Health, Bethesda, MD).

Immunohistochemistry: Sensory ganglia were removed, placed in 4% formalin overnight, transferred to 30% sucrose for cryoprotection for 24 hrs, then mounted in Optimal Cutting Temperature (OCT; #23-730-571; Fisher Scientific) compound. TG and DRG sections were cut into 20 μm slices using a cryostat and mounted onto positively charged (Superfrost plus; #12-550-15; Fisher) slides for immunohistochemistry. Following three 5-minute washes in 1X PBS, the slides were then put into a permeabilization solution containing 10% normal goat serum (NGS; #S13150H; R&D Systems, Mineapolis, MN) and 0.2% Triton X 100 (#9002-93-1; Sigma) in PBS for 30 minutes. This was followed by another series of 5-minute washes in PBS and 1 hr in a blocking solution containing 10% NGS and 0.01% Na azide (#18-613-272; Fisher) in PBS. Following another PBS wash, the slides were incubated overnight in a primary antibody solution made from the blocking solution. The next day, the slides were washed again in PBS then incubated in a secondary antibody solution also made from the blocking solution for 1 hr. Following a PBS wash, counterstaining with DAPI, a PBS wash, and a wash in deionized H_2_O, Prolong Gold mounting media was used to mount coverslips. The primary antibodies used were anti-TRPV1 (1:500; ab203103; Abcam), ATF3 (1:1000; ab207434, Abcam), anti-peripherin, and mouse monoclonal (1:500; P5117, Sigma). Secondary antibodies used were Alexa Fluor anti-rabbit 488, anti-rabbit 568 and anti-mouse 568 (1:1000; Thermofisher). Staining was visualized using a fluorescent microscope (Nikon Eclipse Ni-U) and images were generated using ImageJ.

Nascent protein synthesis “Click Chemistry”: Protein synthesis *in vitro* was measured using Click iT chemistry, as previously established (Melemedjian et al. 2010). TG neurons were isolated, cultured, and treated with sEVs as described above. Following treatment, cells were incubated in methionine-free DMEM (21013024; Gibco) for 30 minutes followed by a 2-hr incubation in 50µM Click iT AHA (L Azidohomoalanine) (C10102; Invitrogen) in methionine-free DMEM. 50 ng/mL nerve growth factor (NGF) treatment was used as a positive control as it induces an increase in protein synthesis (Melemedjian et al. 2010). The cells were then washed with PBS and fixed with cold methanol. This was followed by incubation in 3% bovine serum albumin (BSA; BP9700100; Fisher Scientific) in PBS for 15 min. Cells that incorporated AHA were labeled with Alexa 488–alkyne conjugate (1:200; A10267; Invitrogen) by incubating fixed cells with the conjugate for 30 min. The cells were then washed once with 3% BSA in PBS, followed by another wash with PBS. Cells were then blocked with 10% normal goat serum and labelled with anti-peripherin mouse monoclonal as previously described.

sEVs labeling: Purified sEVs were labeled using the ExoGlow-Protein EV Labeling Kit (EXOGP400A-1, System Biosciences, Palo Alto, CA). sEVs were suspended in 500 µL PBS and 1 µL of the 500X labeling dye was added. This solution was incubated at 37^o^C with shaking for 20 minutes. Following this incubation, 167 μL ExoQuick-TC (EXOTC10A-1, System Biosciences, Palo Alto, CA) was added to the solution followed by an overnight incubation at 4^o^C. The following day, the sEVs solution was centrifuged for 10 min (10,000 rpm), the supernatant was aspirated, and the remaining pellet was resuspended in PBS prior to plantar injection.

Calcium Imaging: Mice *TrpV1*^Cre^:GCaMP6 were used for calcium imaging. TG neurons from these mice were cultured using the previously detailed methods (Inyang et al. 2019). Neurons were treated for 24 hrs with purified sEVs. Coverslips were placed in a custom-designed imaging chamber and and 37°C HEPES-buffered physiological salt solution was superfused across. This solution contains NaCl (134 mM), HEPES (10mM), KCl (6 mM), glucose (7 mM), MgCl_2_ (1.2 mM), and CaCl_2_ (2 mM). Images were recorded using an Andor Zyla 4.2 PLUS scMOS digital camera (Andor, Oxford Instruments, UK) mounted on a variable-zoom Nikon SMZ18 stereomicroscope with 1X SHR Plan Apo objective (NA=0.15). Calcium events were recorded at 10 Hz over the course of 2.5 minutes using µManager software (http://www.micro-manager.org).

Calcium imaging analysis: Calcium oscillations were analyzed using Cell Profiler 4.0 (Stirling et al. 2021). <https://doi.org/10.1186/s12859-021-04344-9>) and GraphPad Prism 9. (GraphPad). Briefly, cells were identified using an image of the tdTomato expression to identify cells that were TrpV1 positive. Then, the integrated intensity of GCaMP6 fluorescence was determined for each TrpV1 positive cell in every frame of the recording. This intensity data was plotted for each cell and the data for cells with a stable baseline and at least 1 calcium oscillation were transferred to GraphPad for analysis. The data were corrected by dividing all values by the baseline and then the area under the curve was calculated for each cell.

Drug administration: Narciclasine (NCLS, sc-361271; Santa Cruz, Dallas, TX), was dissolved in Dimethyl sulfoxide (D128-1; Fisher Scientific) then diluted in 45% w/v 2-Hydroxypropyl-β-cyclodextrin (sc-203461A; Santa Cruz) for i.p. injection at a dose that produces an analgesic effect in preclinical models of persistent pain (Inyang 2019) (Inyang et al. 2019). Rapamycin (1 mg/kg; #37094; Sigma-Aldrich) was dissolved in DMSO then diluted to 10% in saline. GW4869 (1.25 mg/kg i.p.; D1692; Sigma-Aldrich) is a neutral sphingomyelinase inhibitor used for blocking the release of sEVs from the multivesicular bodies (Kosaka et al. 2010, Li et al. 2013, Wang et al. 2014). GW4869 was dissolved to 5 mg/mL in DMSO then diluted to 1.25 mg/kg in PBS. 50 mg QX-314 bromide (#1014; Tocris Bioscience, Bristol, United Kingdom) was dissolved in 1 ml solution (50% saline + 50% pure water) to produce a 5% solution. This solution was diluted in saline to appropriate concentration and injected intraplantar. Ketoprofen (15 mg/kg; K1751; Sigma-Aldrich) was dissolved in 100% ethanol (5 mg in 0.5 mL) then diluted in saline for i.p. injections.

Use of publicly available RNA-seq data and IPA: Human DRG RNA-sequencing data from naïve DRGs were directly downloaded from <https://www.ncbi.nlm.nih.gov/pmc/articles/PMC7305999/> (Wangzhou et al. 2020). Human DRG RNA sequencing data from DRG exposed to human HNSCC-derived sEVs were downloaded from <https://www.ncbi.nlm.nih.gov/geo/query/acc.cgi?acc=GSE134220> and correspond to GSM3939063, GSM3939065, and GSM3939067 (Amit et al. 2020). To normalize the datasets, we matched several hundred genes from the two RNA-seq datasets, and we used the matched housekeeping genes for normalization. We scaled the expression levels of all RNA species in each individual sample. Therefore, the median of ratios between sample and consensus was 1. Genes with missing expression data in one dataset were omitted. To identify gene-enrichment pathways, the data were analyzed through QIAGEN Ingenuity Pathway Analysis (Krämer et al. 2014). QIAGEN Ingenuity Pathway Analysis library identified canonical pathways that were most significant to the data set. The significance of the association between the data set and the canonical pathway was measured in two ways: 1) A ratio of the number of molecules from the data set that map to the pathway divided by the total number of molecules that map to the canonical pathway is displayed; and 2) A right-tailed Fisher’s Exact Test was used to calculate a p-value determining the probability that the association between the genes in the dataset and the canonical pathway is explained by chance alone.

Statistical Analysis: Data are shown as mean ± standard error of the mean and the number of animals or samples used in each analysis are given in figure legends. GraphPad Prism 9 was used to analyze data for statistical tests, which are displayed in figure legends. Repeated measures two-way ANOVAs, One-way ANOVA, unpaired t-test and non-parametric Mann-Whitney tests were used based on experimental design. Statistical significances are indicated as follow * = p < 0.05, ** = p < 0.01, and *** = p < 0.001.
